# Behavioral Phenotyping From deleted CB1 Receptors on Cholinergic Neuron Terminals

**DOI:** 10.1101/2022.06.06.495026

**Authors:** S Wu, K Tsutsui, AY Fitoussi

**Author notes:** Corresponding author: Dr. Aurelie Fitoussi, The Neuroscience Institute, College of Arts & Sciences, Georgia State University, Department of Biology, 444 Natural Sciences Center, 30033 Atlanta, Georgia, United States.

## Abstract

Marijuana is the most widely used illicit drug in the Western Hemisphere and affects physiological processes and cognition. Clear deficits are observed in working memory (WM) that involve the temporary storage and online manipulation of information to solve complex tasks. Marijuana-induced WM deficits have been ascribed to the primary psychoactive compound in marijuana, Δ^9^-tetrahydrocannabinol, which acts at CB1 cannabinoid receptors (CB1r). Recent work emphasized the role of CB1r and cholinergic interaction across this cognitive domain without formal anatomical demonstration. We generated mice with a conditional deletion of CB1r on cholinergic neuron terminals, and WM was evaluated in operant chambers. Control of physiological variables (temperature, nociception, neuromuscular function) was also performed, and additional motor, motivation, time estimation behavior, and effort-based decision-making. Discrete WM enhancement measured in a novel Delay-Non-Matching-To-Position task was evidenced that incorporates early acquisition during randomized delays (mixed procedure), and remarkably, improved performance when these (2s, 8s, 16s, 20s) were kept constant (same procedure) across a testing block of trials. We reported higher motivation in an exponential progressive ratio schedule whilst locomotor activity did not differ between genotypes in the rotarod and open field. However, timing behavior was modified as indicated by higher discriminated motor responses for the shortest interval in conditional deleted mice in the Fixed-Interval time task (10s, 30s). We reported no effect on effort-based decision-making. Our work outlines presynaptic CB1r-cholinergic neuron function(s), and the hippocampus, neocortex, and amygdala brain regions as critical loci through known basal forebrain efferent projections possibly involved in WM and motivation in marijuana intoxication.

## Introduction

Marijuana (*cannabis sativa*) intoxication is a complex phenomenon involving many physiological processes that include tachycardia, hypothermia, and analgesia. These properties are mediated by delta-9-tetrahydrocannabinol (Δ^9^-THC), the (psycho)active constituent of marijuana, which interacts with CB1 receptors in several brain areas [1]. Activation of CB1r is well-known to sustain self-administration of the drug, as well as its pleasurable effects, resulting from its action on the reward circuit [2]. Presynaptic CB1r could centrally induce inhibition of neurotransmitter release via a G-coupled protein also termed depression-induced suppression of inhibition on both excitatory and inhibitory neurons such as the ɣ-aminobutyric acid (GABA), glutamate, and acetylcholine, *a priori* not on dopaminergic (DA) terminals [3, 4]. To what extent presynaptic CB1r could shape cholinergic neuronal function is not entirely known. Short-term memory problems are among the most frequently (additional) self-reported consequences of marijuana use and have been linked to cholinergic system activity. Specifically, temporary information encoding appears to be dramatically impaired [5, 6]. Both endogenous and exogenous cannabinoid administration impaired working memory (WM) [7]. Co-infusion of a CB1r antagonist reversed cannabinoid-induced WM deficits [8]. More importantly, blocking CB1r alone facilitates subsequent WM performance [7]. These effects are thought to mostly arise from disruption of CB1r tone in the hippocampus (HPC) [8], where CB1r are highly expressed and modulate neuronal activity through cholinergic transmission [9] and massive innervation originating from the medial septum [10]. This region is a part of the Basal Forebrain (BF) set of nuclei and constitutes the main source of cholinergic neurons (output), together with the brainstem [11, 12]. BF sends additional important direct efferent projections with presynaptic CB1r to the neocortex and the amygdala, indirectly the striatum, and plays a considerable role in attention, and flexibility [12], and a possible involvement in modulating emotional and motivational processes as suggested by recent work. However, the functional role of CB1r located on cholinergic neurons in this framework is not yet well-characterized.

Here, we generated mice with a conditional deletion of CB1r on cholinergic terminals by first crossing CB1 floxed to mice expressing *Cre* recombinase in cholinergic neurons, thus resulting in mice lacking CB1r on cholinergic neurons (terminals). Animals were tested in several tasks including WM evaluation. The latter is usually based on the retrieval of information across several periods of storage. In the Delay-Non-Matching-To-Position task (D-NMTP) operant schedule, one of two retractable levers is extended as a sample. After a delay period, both levers are extended and the animal has to choose the non-matching lever for reward receipt. Different delay durations, from 0 to 20s, have been tested. Randomized delays presentation across trials (mixed delays procedure) and fixed delays per block of trials (same procedure) were performed. Additional tests including spontaneous alternation in a Y-Maze, interval timing in a fixed-interval time task (2s, 10s, and 30s), primary cost and reward magnitude discrimination in an effort-based choice schedule, and motivation in an exponential progressive ratio schedule were performed, aside from the control of physiological variables including temperature, pain sensitivity in a Hot Plate Test and neuromuscular function in the Wire Hang test, and locomotion in the Rotarod and Open Field tasks.

## Material and Methods

### Animals

Generation of Chatcre-CB1^f/f^ mice. CB1flox (CB1^f/f^) mice express two lox p sites flanking the CB1 receptor (CB1r) gene. Chatcre-CB1^f/f^ were obtained by crossing CB1^f/f^ and ChaTcre+/-mice using a three-step breeding procedure. CB1^f/f^ mice were available from Fisher’s Lane Animal Center (FLAC) maintained by the NIAAA-NIH. Chat-cre lines were available from Dr. Adam Puche’s Laboratory (University of Maryland). All lines were in a predominant C57BL/6N background contribution.

PCR following tail docking was performed to confirm the genotype. Mice were anesthetized with isoflurane and a 4-mm section of the tail tip was obtained. Kwik Stop powder with benzocaine was then applied. Animals were returned to their homecages before the bleeding had stopped. This procedure was performed by the Mouse Consortium directed by Franck Margolis (University of Maryland).

*Housing*. Male mice were used aged from 4 to 8 months. Animals were housed in individual homecages in a temperature-controlled room (22°C) on a 12-hour light/dark cycle (light on at 7:00 AM). Tests were conducted during the light phase of the cycle. They had free access to water and were food-deprived (85% ± 2% of free-feeding weight) throughout the experiments unless stated otherwise. All procedures were conducted in strict accordance with the IACUC protocol (University of Maryland).

### Behavioral tests

*Locomotion* (Open-Field). Animals were free to explore a rectangular white open box for a single 20-min session. Distance and time duration on the center were recorded.

*Locomotion* (Rotarod). A locomotion test was conducted using a cylinder diameter of 31.75 cm from IITC Life Science. When ready to start testing, the animal was placed onto the non-rotating rotarod cylinder. Three testing days were performed including three trials a day. Each trial (ranging from 4 to 40 RPM) lasted 5-min (1 min inter-trial time).

*Catalepsy* (Ring Stand test). Mice were positioned on an 8 cm-diameter ring stand (height 16 cm). The time the animal was motionless was recorded in a 5-min test session. Mice that either fell or actively jumped from the ring were allowed five such escapes.

*Neuromuscular function* (Wire Hang Test). This test was conducted as followed: the mouse was placed on a wire cage lid which was gently waved in the air so that the mouse was able to grip the wire. The lid was then turned upside down, approximately 15 cm above the surface of the soft bedding material. The latency to fall onto the bedding was recorded, with a 60s cut-off time.

*Body temperature*. The body temperature (° Celsus) was measured before the Hot Plate Test. *Nociception* (Hot Plate Test). Nociception function (analgesia) was measured using a Hot Plate analgesia meter. The plate was heated to 55°C ± 0.5°C. The time for the animal to lick its forepaw or hindpaw was recorded. A cut-off time of 30s was set to avoid tissue damage.

*Spontaneous alternation* (in a Y-Maze). Each mouse was placed onto the same starting arm and allowed to freely explore the maze within an 8-min session (from [13]). The number of visits and time spent within the three arms were also recorded.

*Working memory* (Delay Non-Matching-To-Position, D-NMTP). *Apparatus*. Eight identical operant chambers (21.6 cm× 17.8 cm×14 cm; Med Associates, St Albans, VT, USA) housed within sound-attenuating enclosures were used. Each chamber was equipped with two retractable levers (located 2 cm above the floor) and one LED stimulus light located above each lever (4.6 cm above the lever). An external food magazine was connected to a dispenser, centrally located between the two levers, that delivered chocolate-flavored pellets (45 mg, Bio-Serv, Frenchtown, NJ, USA). A houselight, as well as a white-noise speaker (60-80 dB, masking noise background), were located on the opposite wall.

*Protocol*. Procedures were adapted and modified from previous studies (**figure 1**) [14, 15]. Operant training and acquisition of working memory (rule) consisted of several steps. Animals were first trained in a fixed ratio 1 (FR-1) schedule and each lever press led to a single food pellet delivery. The criterion was 60 pellets or 40 minutes whichever came first. After eight sessions, mice were trained in an FR-1 random schedule, and both left and right levers were presented randomly (with the associated top cue-light) so that animals could selectively alternate both sides. The criterion was similar as compared to the previous step. After six sessions, animals were trained in the Easy-Sample step wherein sample and non-matching, choice levers were introduced. Sample lever presentation was randomly alternated between the right and the left side and signaled by the cue light above it. There was no delay between the sample and choice phases, and no punishments. However, an inter-trial period (ITI) of 5 seconds signaled to the animal by the houselight turned off, already separated each trial. Failure in responding to the sample or non-matching lever within 10s resulted in lever retractation and was counted as an omission trial. The total number of lever responses was counted, and the number of correct responses leading to a single food pellet delivery was scored. Accuracy was defined by the percentage of correct lever responses among total lever responses. A stable 80% correct performance validated this stage. The Final-Sample schedule was designated in facilitating working memory non-matching rule acquisition. The incorrect response led to a time-out period of 5s with the houselight turned off, in addition to the non-delivery of the reward. After reaching 80% correct responses (criterion performance), animals were required to make a nose-poke between the sample and choice levers under the Non-Matching-To-Position (NMTP) schedule. In this schedule, after pressing the sample lever within 10s which was then immediately retracted, a nose-poke performed in the back of the operant chamber allowed the presentation of both levers. Animals had to choose the non-matching lever to collect the reward in the food magazine. Priming animal response was necessary early in the schedule. The session ended after 8o trials or 40 min whichever came first. A criterion of 75% correct responses for at least three consecutive sessions was defined. After reaching a stable high level of performance, ITI duration was modified from 5s to 10s, until reaching the same previous criterion. Failure to make a nose-poke after sample lever press within 10s or to the non-matching lever (after presentation) within the same aforementioned time duration was counted as an omission trial. Well-trained animals made a nose-poke immediately after pressing the sample lever. Animals were then evaluated in the Delay-Non-Matching-To-Position (D-NMTP – with 0s) procedure (**figure 1**). This progressive (increasing delays) phase consisted in introducing all the delay duration, with additional instrumental parameters similar to previously. Importantly, 0s was introduced to internally validate the overall schedule. This methodology was applied to make the animal learn to nose-poke consistently during all the delay periods. To this end, this protocol was applied and refined from previous studies (see. [14], [15]):

> – 2 sessions in DNMTP (0-4s): 0s, 1s, 2s, 4s
>
> – 2 sessions in DNMTP (0-6s): 0s, 2s, 4, 6s
>
> – 4 sessions in DNMTP (0-8s): 0s, 2s, 4s, 6s, 8s
>
> – 9 sessions in DNMTP (0-12s): 0s, 2s, 6s, 8s, 12s
>
> – 6 sessions in DNMTP (0-16s): 0s, 2s, 8s, 12s, 16s
>
> – 6 sessions in DNMTP (0-20s): 0s, 2s, 8s, 12s, 20s

**Figure 1.**
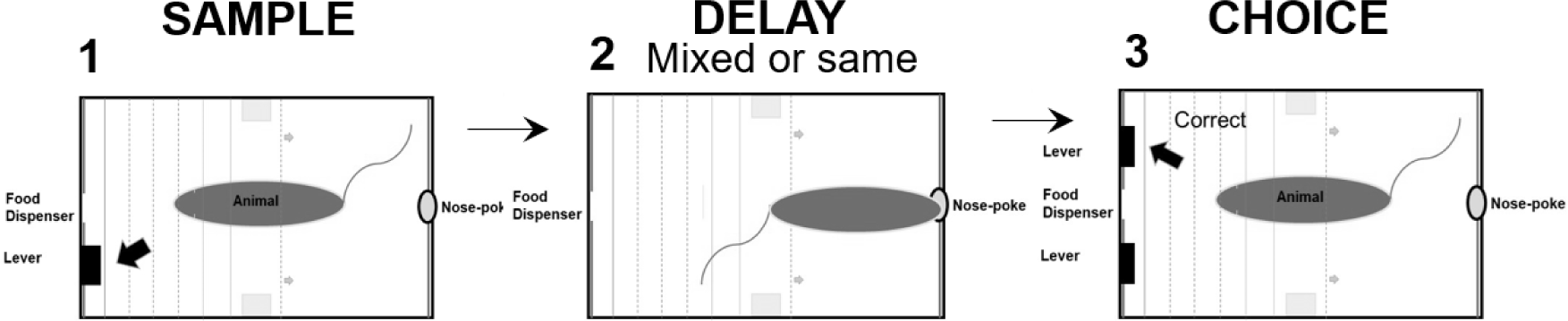
Delay-Non-Matching-To-Position task. Right or left levers are randomly presented during the sample phase. After pressing the sample lever, a delay phase occurs. In either the mixed or same procedure, the animal nose-poke constantly during the delays, presented either randomly (mixed procedure) or subsequently and constant per block of trials (same procedure). The nose-poke at the end of the delay period triggers lever extension. The animal has to choose the opposite lever (non-matching lever) during the choice phase to validate a correct trial.

Delays were randomly introduced during sessions i.e., the *mixed* delays procedure. The latency between the end of the delay and the first nose-poke leading to lever extension was recorded. Next, mice were tested in the *same* delay (DNMTP – 0s missing) procedure in which only 2s, 8s, and either 16 or 20s were evaluated sequentially and presented per block of trials, thirty trials per delay condition.

*Motivation* (exponential progressive ratio schedule). In well-trained mice, animals were additionally trained with one session of FR-1 and two sessions of FR-5 (5 lever presses resulted in reward delivery) before the exponential progressive ratio (PR) schedule. Under PR, the response requirement on the active lever (set in a counterbalanced fashion) increased trial by trial exponentially as described previously (see. [16]) to earn a reward. After reward delivery, levers were retracted for 2s before the onset of the next trial, and the houselight was turned off during this inter-trial time. The maximum number of lever presses provided by the animal through trials is called the « breakpoint » and was used as a motivational index. For a detailed sequence of lever ratio implementation, see. [16]. The second batch of naive animals underwent an operant training schedule as previously published and tested in PR as described above.

*Temporally control of behavior*, i.e., timing (Fixed interval time schedule, FI). In well-trained mice, animals were additionally trained with one session of FR-1 and two sessions of Fixed Interval 2s (FI-2s) schedule. Under this schedule, trial onset was signaled to the animal by lever extension, and this was also associated with the starting interval duration (i.e., 2s). Responding within the interval, in either the active or inactive lever had no instrumental effect (i.e., no food reward). However, the first response made on the active lever after the end of the interval resulted in food reward delivery, followed by a 10s-lever retraction period. After two sessions and stable lever responses provided, animals were switched to FI-10s and FI-30s respectively, under which the interval duration was set from 2s to 10s and 30s. Animals were switched to the FI-10s to FI-30s schedule after reaching stable performance (i.e., six sessions).

*Effort-based decision-making primary ratio.* In well-trained mice, animals were additionally trained with three to five sessions of FR-1, 60 pellets (validated criterion, three consecutive sessions), or 40 minutes whichever came first. Then, mice were trained in a forced choices (FC) schedule wherein the two options that differed in terms of reward magnitude and lever response effort were presented to the animals, randomly and alternating with both left and right sides. Either ten lever presses to earn three pellets or one lever press to earn a single food pellet were available through fifteen trials. The next day, a mixed session with ten forced trials and a subsequent fifteen free trials were achieved. Finally, choice preference between the two options was evaluated during twenty-five free trials, and until stable preference for at least two consecutive sessions was demonstrated (stability was defined as <15% variation between sessions).

## Data analysis

Independent t-test or One/two-way and repeated measures ANOVAs, when required, were performed for genotype comparisons using dedicated behavioral parameters as mean ± sem and using Statistica software 10. F value (group factor) was indicated (significance threshold, 5%, p_value_ < 0.05). Post-hoc analysis completed variance analysis (for p_value_ < 0.05) using PLSD Fisher’s test (significance threshold, 5%, p_value_ < 0.05).

### Results Open-field

This test (n = 15) that examines exploration patterns revealed no statistical difference in the total distance traveled (*t-test*, t = 1.10, ns), as well as within the session when examining 5-bin minute periods (**figure 2A**). The percentage of time mirrored the distance parameter (data not shown). Finally, time spent in the center was stable during the session, and similar between both groups (control: n = 5; experimental: n = 10; *t-test*, t = 1.25, ns).

**Figure 2.**
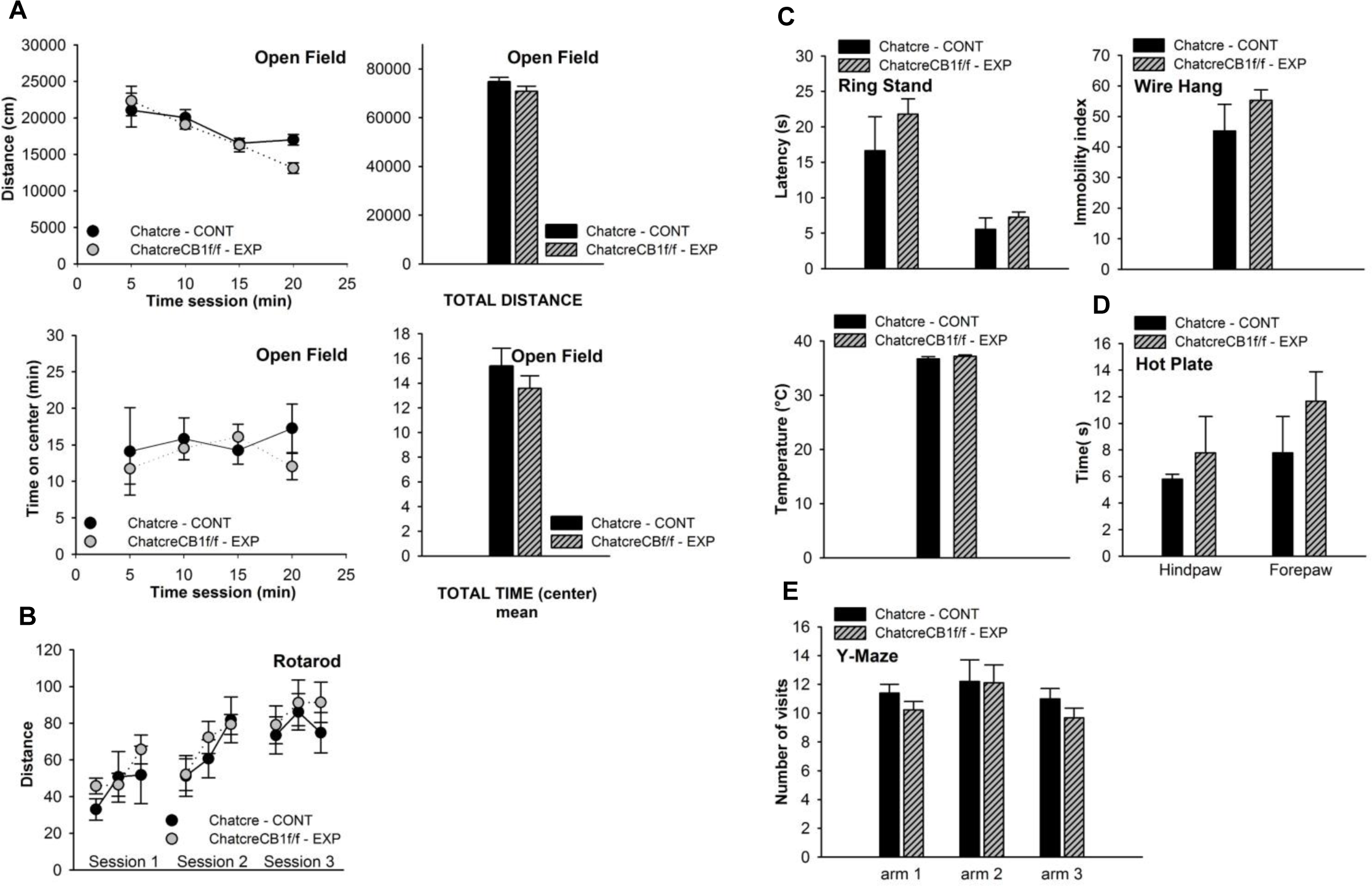
Preserved physiological parameters in ChatcreCB1^f/f^ mice (experimental, n = 10) and CB^f/f^ (control, n = 5). (A) Locomotor function evaluated in the open-field (OF) and (B) rotarod. Animals displayed preserved locomotor activity in OF i.e., distance traveled (time-course), total distance, time on center (time-course), total time on center, and preserved locomotor ability evaluated in the rotarod as indicated with the distance traveled through 3 sessions, 3 trials a day, (C) Catalepsy, neuromuscular function, and temperature. The Ring Stand test consisted in evaluating latency and immobility, similar between genotypes. The Wire Hang test consisted in measuring the latency to fall and revealed preserved motor function. Animals displayed preserved temperature (° Celsius), (D) Nociception. Animals displayed similar times to lick hinds- and forepaws when heated in the Hot Plate, and (E) Spontaneous alternation was also preserved as measured in the one-session Y-maze and the number of arm visits within the 3 available arms. *ANOVA, no significant effect was found*.

### Rotarod

Locomotor activity in this task (n = 15) was recorded for three consecutive days and three trials a day (**figure 2B**). Overall, increased locomotor activity was revealed as illustrated when comparing the first to the third day (*day*, F_1,15_ = 16.65, p<0.001), but no significant group difference (control: n = 5; experimental: n = 10) was observed (*group*, F_1,15_ = 3.21, ns).

### Ring Stand and Wire Hang Tests

Both tests (n = 15) failed to reveal significant differences from CB1r deletion on cholinergic neurons. As illustrated in **figure 2C**, latency to jump was scored 16.64 ± 4.79 for control (n = 5) and 21.79 ± 2.17 for genetically-modified mice (n = 10) in the Ring Stand Test (*t-test*, t = - 1.13, ns), and also, see. immobility index in **figure 2C** (*t-test*, t = - 1.13, ns). On the Wire Hang Test (*t-test*, t = −1.30, ns), latency was scored 45.25 ± 8.75 for the control and 55.24 ± 3.46 for the conditional knockout mice.

### Temperature

Conditional knockout (n = 5) and control mice (n = 10) had the same temperature (*t-test*, t = - 1.13, ns), with 36.72 ± 0.39 and 37.19 ± 0.22 respectively (**figure 2C**) indicating no significant alteration from the conditional CB1r deletion.

### Hot Plate test

We found that both groups (n = 15) of mice were sensitive to heat and licked within the same time interval, for both forepaw (*t-test*, t = −0.53, ns) and hindpaw (*t-test*, t = −0.74, ns), as shown in **figure 2D** (control: n = 5; experimental: n = 10).

### Y-Maze

This one session measure (n = 15) of spontaneous alternation and exploration revealed that animals displayed the same level of exploratory behavior when examining the percentage of time spent in the three arms (*t-test*, t = 1.33, ns) as compared to the session duration (*t-test*, t =

-0.17, ns) (data not shown). Detailed analysis failed to extract relevant differences in terms of arm visits (*t-test*, t = 0.76, ns) (**figure 2E**), or time spent in these arms. For instance, and as shown in **figure 2E**, the number of visits in arm 1 (right arm) was scored 11.40 ± 0.60 in control (n = 5) and 10.22 ± 0.60 in experimental (n = 10) mice. Distance traveled in arm 1 and adjacent arms did not reveal statistical difference (*t-test*, t = 1.07, ns) as indicated in this arm: 5571 mm ± 220 in control and 5582 mm ± 203 in experimental; arm 2: 6588 ± 402 in control and 6419 ± 608 in experimental and arm 3: 6275 ± 442 in control and 5205 ± 298 in experimental.

### Working memory

This study (control: n = 7; experimental: n = 6) emphasized the effect of the conditional CB1r deletion on working memory capacities. First, animals were trained in a fixed ratio 1 and fixed ratio 1, random, as illustrated in **figure 3A**. They all acquired this short operant training (*t-test*, t = −0.74, ns; FR1 rand: *t-test*, t = −1.10, ns). Animals performed the Easy Sample step in which they had to alternate their behavioral response asked by the random alternation of lever presentation to obtain a single food pellet. Although no time-out (as a penalty) indicated to the animal that a wrong response was performed, the non-matching rule was already introduced at this step with no delays and no nose-pokes between the sample and choice phases (**figure 3B**). Both groups acquired the rule and reached more than 80% correct responses (*t-test*, t = 1.13, ns). After these sessions, they were on the Final SA schedule (**figure 3C**) that consisted, essentially, of adding a time-out period (5s) when an incorrect response was provided. After reaching, similarly, 80% of correct responses, animals were evaluated in the Non-Matching-To-Position schedule (NMTP) in which making a nose-poke was required between the sample and choice phases (no delays at this step, **figure 3D**). Extension of the levers was not possible until the animal had made successfully a nose-poke. After reaching equally a significantly high number of correct responses, i.e., more than 70% (*t-test*, t = 1.07, ns), animals underwent the specific protocol of progressive delay implementation in the Delay-Non-Matching-To-Position, D-NMTP (0s, 2s, 4s, 6s, 8s, 12s, 16s, 20s) (**figure 4**) (see. Material and Methods section). On this occasion, we revealed significant differences in working memory acquisition (*group*, F_1,13_ = 9.97, p<0.05), that persisted, sometimes sporadically, for some delay duration, and no significant improvement for 16s (post-hoc, ns) and 20s (post-hoc, ns). But most of the time, animals reached the same level of performance, see. the three last sessions 0s (post-hoc, ns), 4s (post-hoc, ns), 12s (post-hoc, ns), 16s (post-hoc, ns), and 20s (post-hoc, ns) suggesting an improvement in working memory acquisition, rather than performance *per se* (**figure 4**). Specifically, the longest delay (16s and 20s) durations were found to mask genotype differences (16s: *group*, F_1,13_ = 5.67, ns; 20s: *group*, F_1,13_ = 7.10, ns) unlike short and mid-delay durations (*delay*, F_1,13_ = 9.18, p<0.05) i.e., 2s (p<0.05), 4s (p<0.05), 6s (p<0.05) and mostly, 8s (p<0.05) during this mixed delays procedure, when delays were presented randomly across sessions; and when animals had to nose-poke during the whole delay duration, with one nose-poke necessary at the end of the delay period to induce lever extension. During this progressive operant schedule, correct responses (**figure 5A**) were higher in the conditional knockout mice (*group*, F_1,13_ = 9.80, p<0.05) unlike the total number of nose-pokes performed during the delays (*group*, F_1,13_ = 1.30, ns). The latter augmented significantly throughout the progressive schedule implementation, see. session 1 (>50) *vs*. session 30 (>1000) (**figure 5A**). Mean latency between the end of the delay period and the first nose-poke leading to lever extension was scored inferior to 2s early in this training schedule (two first sessions), whereas inferior to 1s late in the procedure (two last sessions) (data not show, *group*, F_1,13_ = 6.57, ns). Clear improvement (*general group means correct responses*, F_1,13_ = 11.65, p<0.05 and *interaction* group × day, similar) as compared to the control was then, revealed during the same delays procedure (**figure 5B**), in which the same delay was kept constant through a block of trials, to lower the cognitive demand and the complexity of the task. Only three delays were presented across sessions, thirty trials per delay condition. Mean latency between the end of the delay period and the first nose-poke to extend levers was displayed in **Table 1**: values tend to decrease throughout sessions and reached about half a second for both groups. In such conditions, improvement in the ChatcreCB1^f/f^ mice was reported at 2s (*group*, F_1,13_ = 8.89, p<0.05), 8s (*group*, F_1,13_ = 16.97, p<0.05), 16s (*group*, F_1,13_ = 15.30, p = 0.07) and 20s (*group*, F_1,13_ = 13.57, p<0.05).

**Figure 3.**
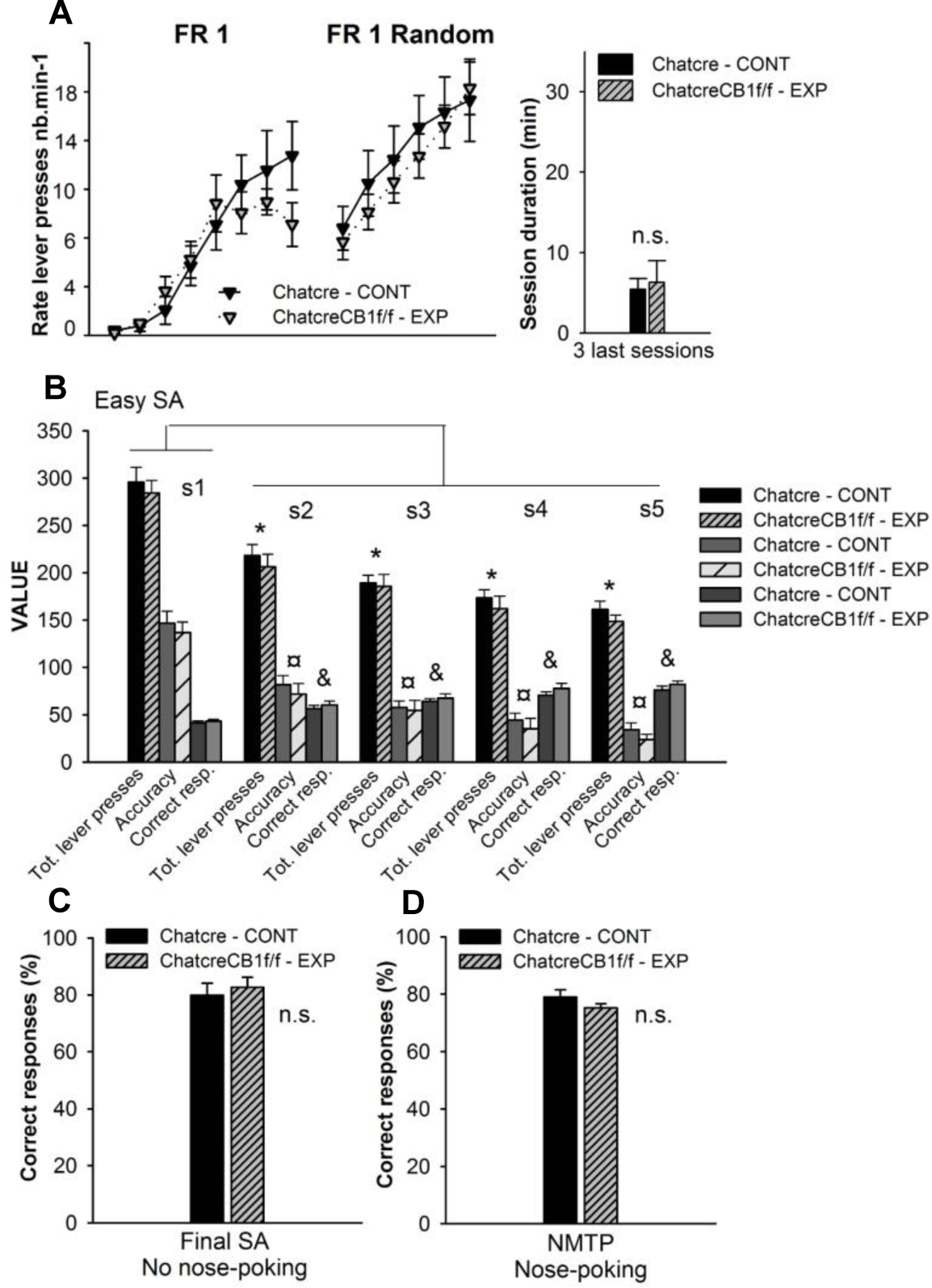
Operant training preceding working memory assessment. No genotype difference (experimental: n = 7; control: n = 6) was reported in either (A) fixed ratio 1 and a random version of this step, supported by the session duration (see. mean three last sessions). Acquisition of the non-matching lever rule started at the next step, the Easy SA (Sample Alternation) (B) and animals progressively decreased total lever presses, improved accuracy, and increased correct responses; (C) Correct responses, about 80% was obtained in the Final SA schedule wherein incorrect response led to a 5s time-out period, and 10s inter-trial time, and (D) Non-Matching-To-Position schedule (NMTP) when a nose-poke was required to extend levers after sample lever presentation. *ANOVA, no significant group difference was found, but session factor was found significant during the Easy SA step, * p<0.05 (total lever presses), ¤ p<0.05 (accuracy), & p<0.05 (correct responses)*.

**Figure 4.**
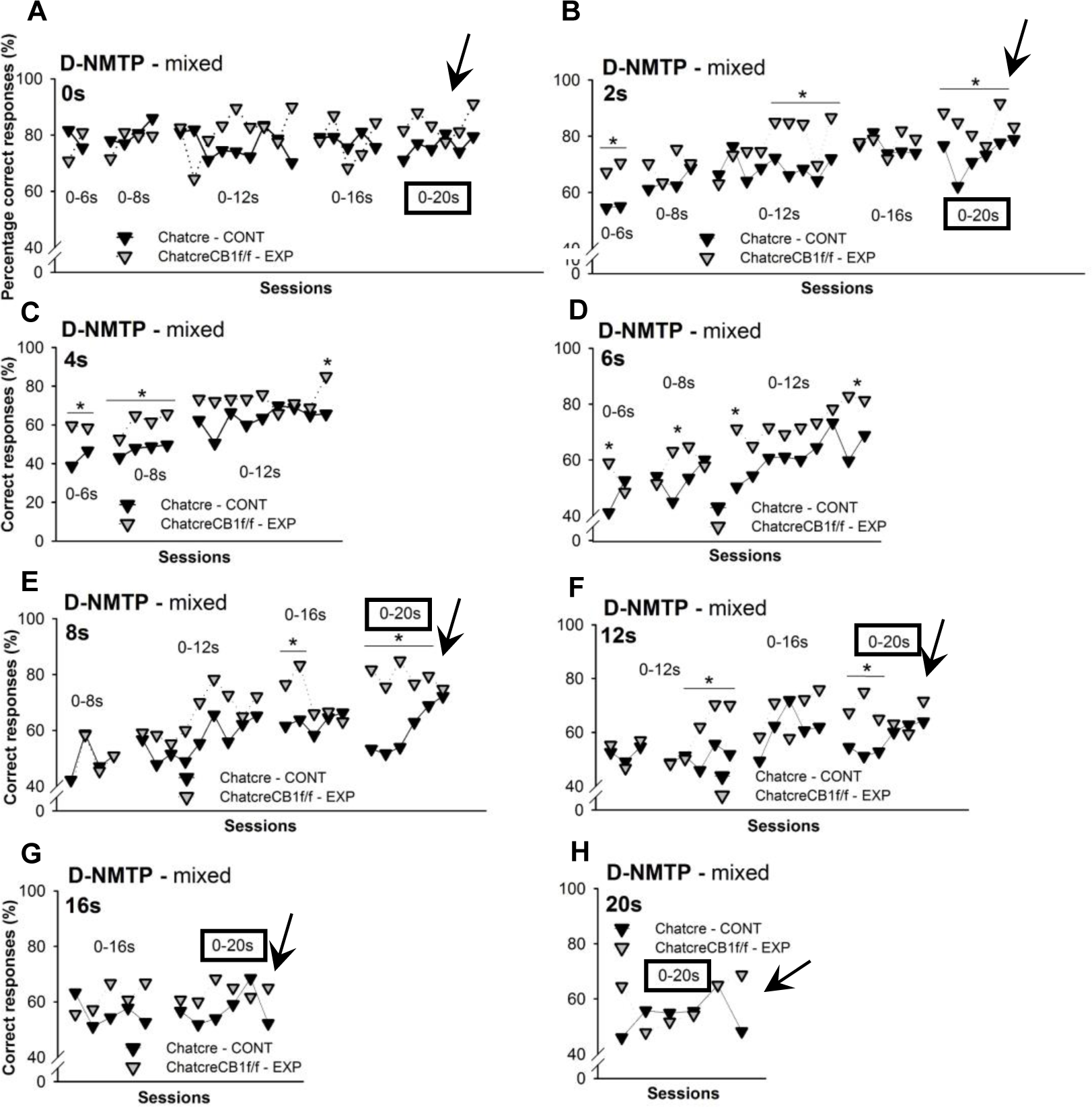
Higher performance in Delay-Non-Matching-To-Position procedure with mixed delays in genetically-modified animals. Varied delay durations were assessed and progressively implemented through the procedure, with the percentage of correct responses at (A) 0s, (B) 2s, (C) 4s, (D) 6s, (E) 8s, (F) 12s, (G) 16s, and (H) 20s displayed. WM assessment was validated for the 3 last sessions of schedules 0-20s in experimental (n = 7) and control (n = 6) mice. Otherwise, responses performed were considered as acquisition only. *ANOVA, * p<0.05.* The black frame signaled the test session during the 0-20s schedule with the black arrow showing the specific testing sessions.

**Figure 5.**
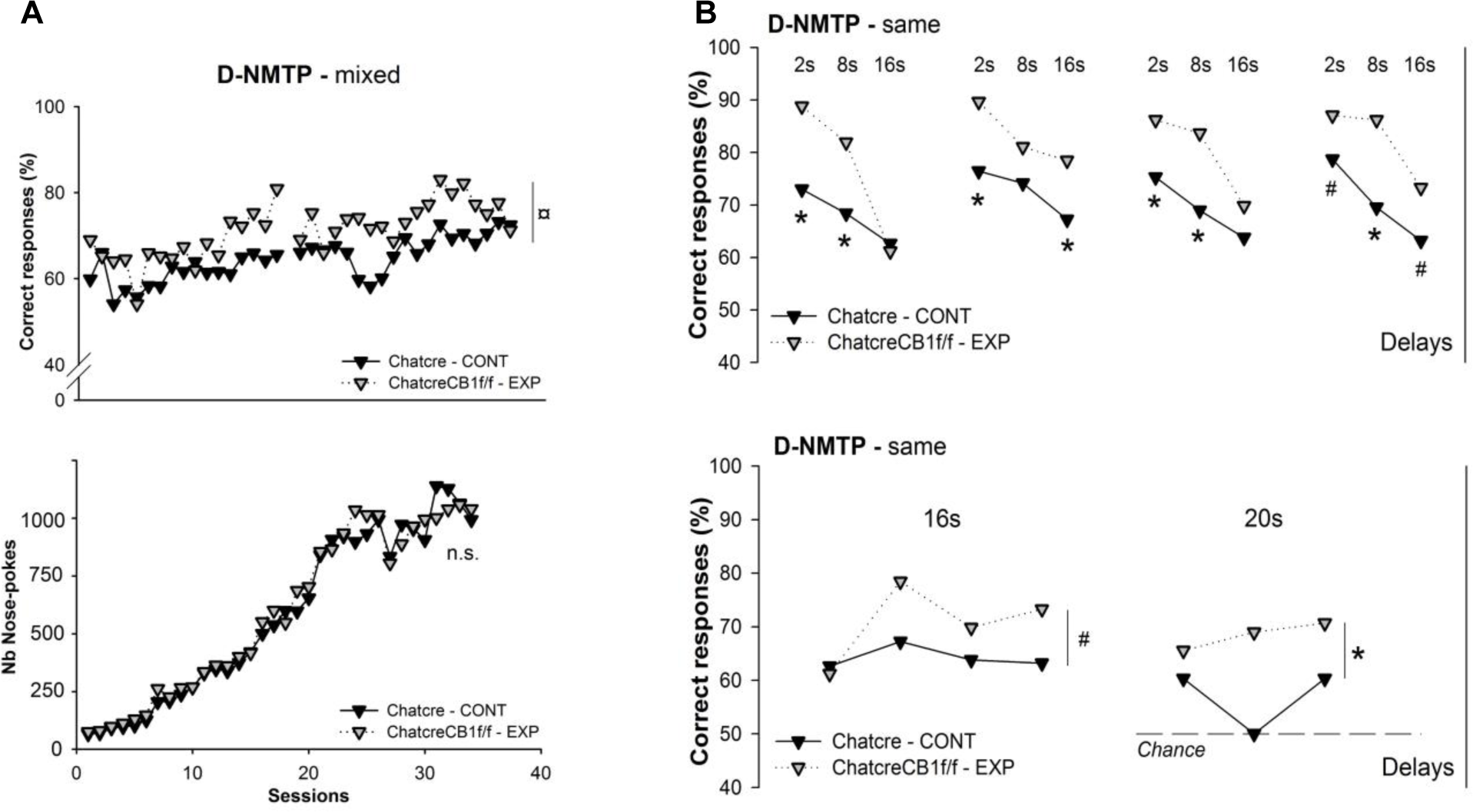
Discrete WM enhancement during the Delay-Non-Matching-To-Position procedure with mixed delays and same delays. (A) Progressive increase in the correct number of responses per behavioral session revealed a discrete acquisition improvement in the D-NMTP for the experimental (n = 7) versus control (n = 6) group, unlike the total number of nose-pokes made during the delay periods; (B) the same delays procedure revealed improvement in the experimental group at 2s, 8s, 16s, 20s as compared to control. *ANOVA, * p<0.05, # p=0.07*.

**Table 1.** Average latency recorded during the same procedure, WM evaluation. The period between the end of the delay duration and the first nose-poke response was displayed and showed a decrease in latency throughout sessions with no genotype difference. This parameter is thought to reflect learning of the task (non-matching rule) independently of performance during the task (DNMTP).

| Same. | **s1** |  | **s2** |  | **s3** |  | **s4** |  |
| --- | --- | --- | --- | --- | --- | --- | --- | --- |
| EXPERIMENTAL | 0,94 | ± 0,23 | 0,56 | ± 0,13 | 0,50 | ± 0,075 | 0,40 | ± 0,08 |
| CONTROL | 0,82 | ± 0,24 | 0,70 | ± 0,14 | 0,46 | ± 0,11 | 0,57 | ± 0,05 |

### Motivation

We found that ChatcreCB1r^f/f^ mice displayed higher lever presses (*group*, F_1,19_ = 11.15, p<0.05; *interaction* group × session, similar), and breakpoint (BP) (*group*, F_1,19_ = 13.45, p<0.05 and *interaction* group × session, similar) in the exponential PR task as measured throughout six sessions and reaching a stable behavior. Interestingly, either a progressive operant training under fixed-ratio schedules or consecutive fixed ratio 1 with 10s ITI (data not shown) led to such higher PR performance in the conditional knockout mice indicating that the operant assessment of motivation was poorly dependent on the operant training and strengthened the effect of the conditional deletion of CB1r. Mean BP value could be approximated to 800 for genetically-modified mice (n = 8) and 600 for control (n = 11) (**figure 6A**). Behavioral responses on the active lever mirrored the decrease in the number of lever responses on the inactive lever (**figure 6A**), and no significant difference between both genotypes (see. session 1 and session 6) (*group*, F_1,19_ = 1.14, ns).

**Figure 6.**
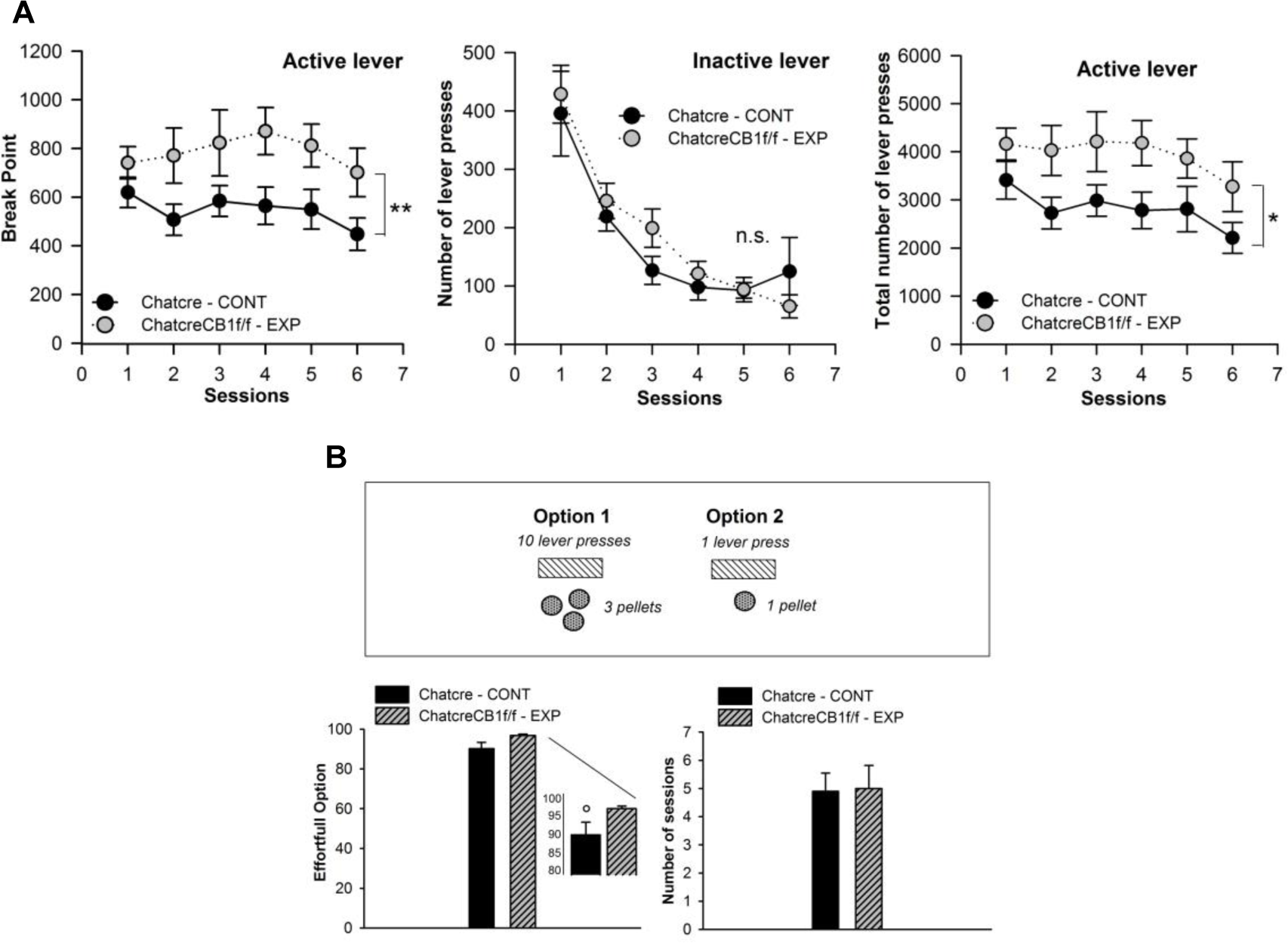
Exponential progressive ratio and effort-based choice schedule. (A) Higher motivation (experimental, n = 8; control, n = 11) as revealed by the higher breakpoint, and a total number of lever presses unlike the decline of instrumental responses on the inactive lever, and (B) Both genotypes (experimental, n = 6; control, n = 10) chose preferentially the high effort-magnitude ratio option, i.e., 10 lever presses for 3 pellets against 1 lever press for 1 pellet in an effort-based choice paradigm. *ANOVA, ** p<0.01, * p<0.05*.

### Effort-based decision-making

When animals (control: n = 10; experimental: n = 6) had to choose between either three pellets but ten lever presses to obtain such a reward, and one pellet but one lever press, all groups chose the high effort but high magnitude option (*t-test*, t = 1.10, ns). The same number of sessions (4 < *n* < 5) was recorded so that all animals reached stable performance. Here, more than 90% of choices were directed toward the high effort-high magnitude option, and we found no statistical difference between the final performance level reached (**figure 6B**).

### Fixed-Interval time task

Animals (control: n = 10; experimental: n = 8) were tested for a fixed interval time task in which 2s, 10s, or 30s interval rule failed to lead to food pellet delivery if pressing the lever during the interval (**figure 7 and figure 8**). Such outcomes could occur after the first lever is pressed at the end of the designated interval. Lever presses through the interval were however recorded, and the active lever (left or right side) was assigned in a counterbalanced fashion. We found no significant differences during the FI-2s for the total number of lever presses (*t-test*, t = −0.10, ns). Interestingly, the pattern of lever presses during FI-10s was modified across the six evaluated sessions (**figure 7B**). The third 2s-bin period was particularly sensitive and a motor shift was observed so that the highest lever responses were provided during the last 2s-bin interval (**figure 7C**). This was more evidenced when examining the non-cumulative lever responses through the response frequency parameters. No significant difference was reported when examining the total number of lever presses (*group*, F_1,18_ = 2.58, ns), or the total number on the inactive lever (*group*, F_1,18_ = 3.69, ns) unlike the non-cumulated response frequency (group, F_1,18_ = 12.54, p<0.05). When normalizing overall locomotor activity and expressing each time epoch as the percentage of the total locomotor activity, the response curve from both genotypes could be superposable. On FI-30s, all parameters were similar (total lever presses: *group*, F_1,18_ = 16.87, ns; lever presses inactive: *group*, F_1,18_ = 14.78, ns; non-cumulated response frequency: *group*, F_1,18_ = 2.54, ns) (**figure 8A**). All animals acquired and expressed similar FI-30s rule i.e., preserved timing behavior. Animals increased progressively and accurately the number of lever presses to obtain the reward, with the highest number of responses provided during the two last 6s-bin of interval achievement duration, similarly for both genotypes (**figure 8B**).

**Figure 7.**
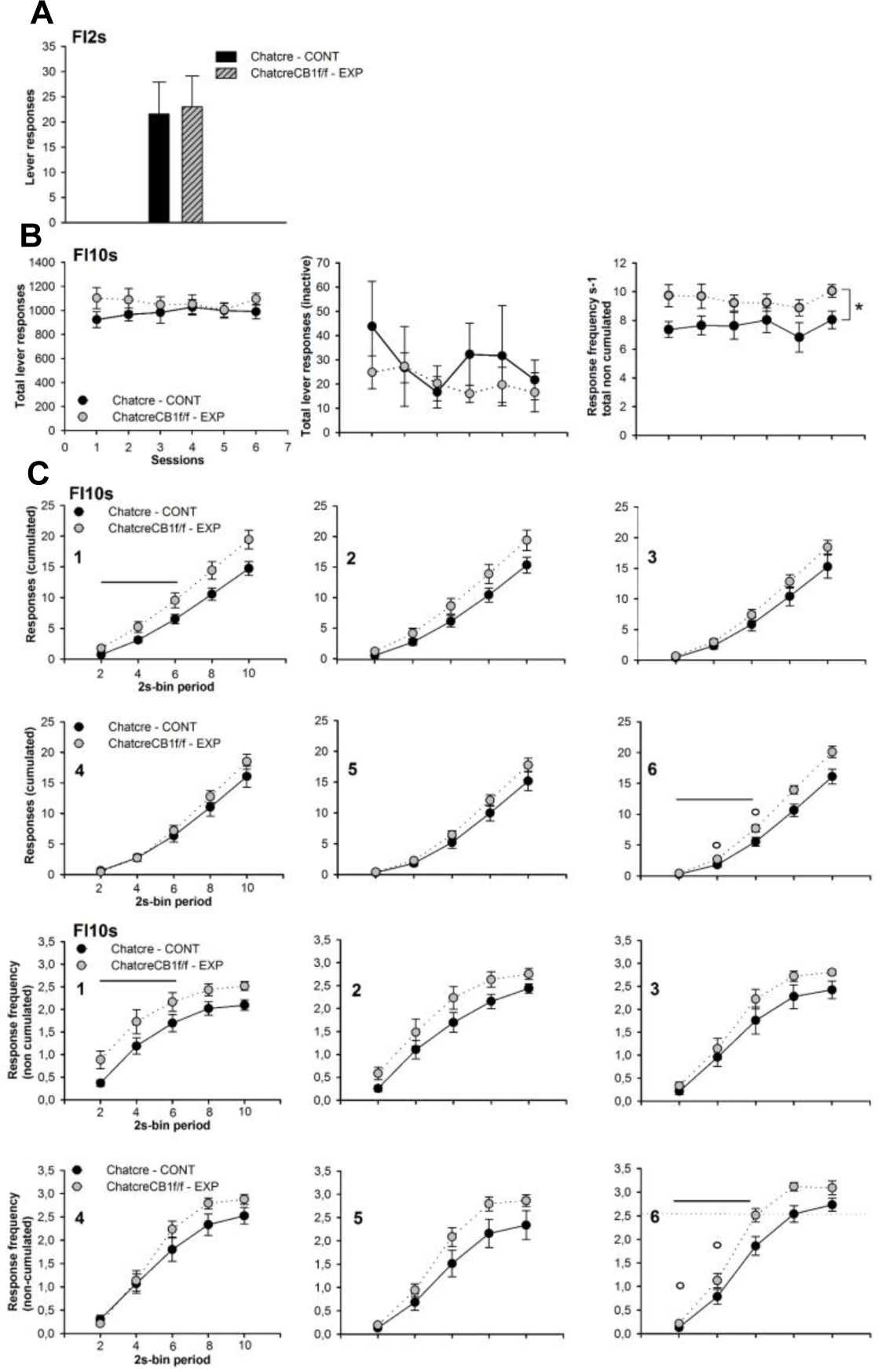
Fixed-interval time task (2s and 10s). (A) 2s-interval lever presses, (B) 10s-interval with total lever presses, and lever responses on the inactive lever revealed a similar pattern of behavioral responses (experimental, n = 8; control, n = 10) whereas response frequency (non-cumulated) in seconds demonstrated higher lever responses for the genetically-modified mice. The detailed pattern of lever responses throughout the 10s-interval was shown with both (C) cumulated responses and non-cumulated responses frequency. Shifting responses early in the interval is thought to reflect the learning rule. *This symbol ° represents the previous behavioral score (session 1). ANOVA, * p<0.05*.

**Figure 8.**
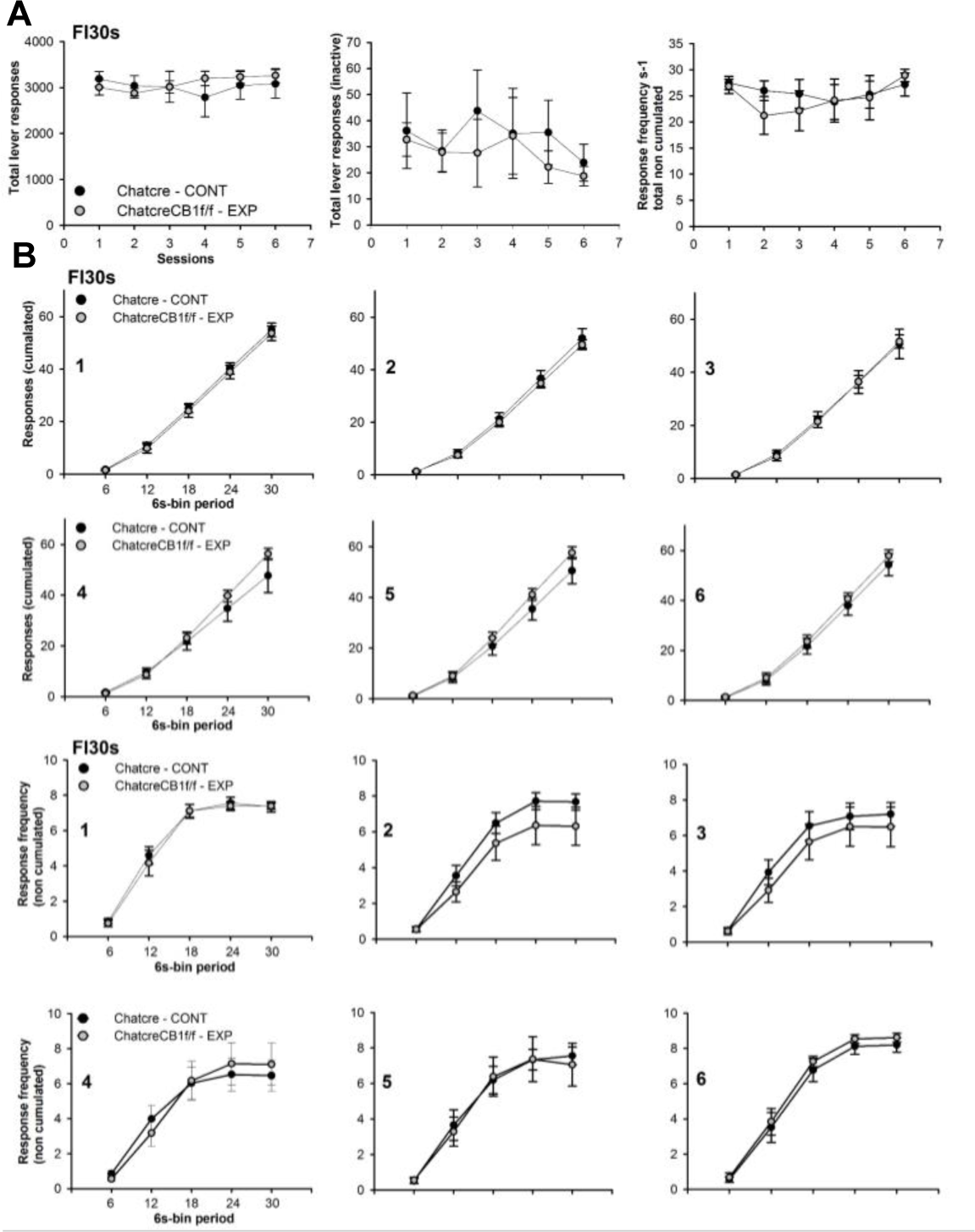
Fixed-Interval time task (30s). (A) 30s-interval with total lever presses, lever responses on the inactive lever, and response frequency revealed a similar pattern of behavioral responses. The detailed pattern of lever responses throughout the 30s-interval was shown with both (B) cumulated responses, and non-cumulated responses (experimental, n = 8: control, n = 10). *This symbol ° represents the previous behavioral score (session 1). ANOVA, No effect*.

## Discussion

In this study, we aimed at characterizing the role of CB1r specifically localized on cholinergic neuron terminals that are mostly represented by the efferent acetylcholinergic projection terminals from the BF and brainstem nuclei [11, 12, 17]. To achieve this goal, we generated mice that display conditional deletion of CB1r on cholinergic neurons (on terminals, see. [18]) with *Cre* enzyme expressed specifically on these cholinergic neuronal populations. Additionally for producing conditional deletion of CB1r in the rat brain, *Cre* expression will allow brain circuit manipulations in targeted neurons with, for instance, light-sensitive channel protein expression, and activation [19], in future relevant work.

We found selective differences in WM abilities as a function of enhanced delay duration. An exception however occurred at the longest delay(s) suggesting that CB1r-dependent cholinergic transmission poorly improves performance at the highest holding period duration along with the complexity of the task. The mixed delays procedure illustrates the predominant facilitation of WM acquisition in genetically-modified mice, whereas the same delays procedure demonstrated a discrete enhancement of WM capacity in the same mice. Additional facilitation in lever responding during working memory acquisition under D-NMTP schedule changes attributable to the conditional deletion was reported. While fundamental physiological variables appeared to be well-preserved i.e., temperature, pain threshold, and neuromuscular function, higher motivation as measured in PR was revealed unlike primary motor abilities indicating that loss of CB1r on cholinergic neurons is not enough to cause drastic motor impairment. However, disparate lower primary motor efficiency in early training could be evidenced in some ChatcreCB1r^f/f^ cohorts (data not shown). Failure to reveal improvement in temporary control of behavior as measured in interval timing completed this preclinical picture, but conditional knockout mice provided higher lever presses toward the active lever at 10s interval (discriminative responses). Finally, we also reported no significant effect of the conditional CB1r deletion in a two choices effort-based schedule.

### Distribution of both presynaptic CB1r and cholinergic projections in the rat brain (involved in cognition)

CB1r are found throughout the brain [20] with the largest expression in the hippocampus, the striatum, and the neocortex [17]. These presynaptic inhibitory receptors inhibit the neurotransmission of both excitatory and inhibitory neurons, including the acetylcholinergic population [21]. This group of neurons (output) is mainly represented by the basal forebrain (BF) and the brainstem set of nuclei [12, 22] that project, essentially with presynaptic CB1r, to the medial septum [23], neocortex, hippocampus (HPC) [24] and the amygdala [11], all of these brain areas predominantly involved in cognition although additional cholinergic projections have been evidenced to the Ventral Tegmental Area and the Thalamus for instance [12]. In this framework, cortical cholinergic projections would constitute a minor component of cholinergic functioning [12].

Presynaptic CB1r at terminal fields have been evidenced using *in situ* hybridization, or autoradiography and immunocytochemistry, outlining the modulation from the endocannabinoid system (eCB) [17, 21]. Additional cholinergic interneurons could be found in the striatum (aspiny CIN), a small proportion directly in the neocortex, and in the HPC from which cell-type identity is in dispute [11]. CIN are unlikely to exhibit presynaptic CB1r although the efficient detection of both acetylcholinergic transferase (Chat) enzyme and CB1r mRNA (i.e., colocalization) could be discussed [17, 25].

### Working memory, attentional processing, and flexibility

The implication of CB1r specifically expressed on cholinergic neurons in WM was expected, but whether this deletion could enhance cognitive performance, presumably through a discrete increase of local ACh tone, remained to be demonstrated. As previously exposed, a large body of evidence showed that stimulation of CB1r with an agonist impaired WM abilities while blocking produced the opposite effects [26], and was directly upon the dependence of HPC neuronal firing [27]. Electrical stimulation of this region reversed the deficits and this is accompanied by changes in neuronal firing [9], through local cholinergic transmission [26]. Intra-HPC blocking with cannabinoid antagonist (i.e., rimonabant) facilitates WM performance while both *in vitro* and *in vivo* cannabinoid agonists application in HPC inhibits ACh release [28] suggesting a predominant implication of CB1r in modulating HPC cholinergic transmission. Further local and systemic CB1r blockade increased HPC ACh levels, possibly through intra-HPC DA-dependent mechanisms but not the genetic deletion [30]. It favored that long-term deletion induces large neurobiological compensations, however, higher ACh HPC levels could be evidenced when the region was highly recruited thus, facilitating subsequent cognitive performance, specifically in learning and memory as demonstrated in CB1r null mutant mice [31] and supported by the behavioral facilitation scored under D-NMTP schedule changes in the early-mid acquisition, and discrete WM enhancement reported in our study. Interestingly, the CB1r agonist applied directly in the medial septum did not affect ACh levels in the HPC [30]. This region provides the main input to the HPC [31] and septal lesions induced short-term memory impairments [32] although contrasting results have been evidenced [33]. This set of data is also consistent with the involvement of HPC and the PFC in flexibility (i.e., adapting behavioral responses in changing environments) [34] and attentional processing [35]. Interestingly, mice overexpressing the vesicular acetylcholine transporter were impaired in short-term WM together with an increase in ACh tone measured with *in vivo* microdialysis [36]. A large array of memory-based deficits was observed unlike motor improvement indicating that a suboptimal increase in ACh level produces detrimental cognitive outcomes and could be comparable with some inefficient cholinergic drugs [37], among cholinergic enhancers specifically [38, 39]. Consequently, improvement in cognitive functions including WM performance favored a discrete increase in ACh tone or cholinergic excitability in ChatcreCB1^f/f^ mice.

Overall, similar spontaneous behavior in Y-maze is consistent with the discrete WM enhancement observed in DNMTP. Although cognitive evaluation in this task does not yet reach a consensus regarding the conceptual framework, it certainly involves exploration and short-term memory (exploration: [40]; short-term memory: [13]; others: [41]) and is not sensitive to age-dependent cognitive decline [41].

The abundance of CB1r was also reported within the prefrontal cortex (PFC) [8], and cannabis exposure induced correlated changes in metabolic activity in this region, and increased Immediate Early Gene expression [7], cannabinoid-induced WM improvement is, however, likely to arise from predominant HPC modulation (as exposed) rather than PFC. Interestingly, regulated feedback could involve the GABAergic neuronal population and the nucleus accumbens [42], a region also known for the emergence and proposed substrate of motivated behaviors [43].

### Temporally-controlled of behavior

We also reported slight modifications in interval timing behavior, which is coherent with the effects of nicotine exposure [44, 45, 46] and cannabinoid drugs [47]. Specific interaction of both systems is firstly evidenced in this study, augmenting overall behavioral responses toward the discriminated reward lever for short intervals (10s) although accuracy *per se* seemed to be preserved. This indicates that CB1r deletion could, eventually, attenuate cholinergic transmission efficiency but did not drastically disturb performances in chronically (i.e., genetically) CB1r deleted animals. Consequently, additional pathways or neurotransmitter systems within the PFC [48] or striatal DA are better predictors of interval timing capacities [49, 50], but indirect modulation of DA could be involved in ChatcreCB1r^f/f^ mice as suggested by previous work.

### Motivation

Interestingly, we outlined higher motivation evaluated in a PR task, independently of primary motor response requirement that points to a role of CB1r in specifically modulating cholinergic transmission for emerging emotional and motivational processes. Preclinical evidence has shown bidirectional crosstalk between nicotinic acetylcholine and eCB systems in brain reward pathways including the limbic system and the prefrontal cortex [3, 51]. This is particularly demonstrated in the effects of eCB on nicotine addiction, and the nicotinic acetylcholinergic system on cannabinoid dependence [52, 53]. For instance, the rewarding effect of nicotine is blunted in CB1 null mutant mice and CB1r activation increased the motivation to self-administer nicotine as measured in the PR task [54]. Additionally, CB1r antagonism dose-dependently decreases nicotine self-administration [55] whilst chronic treatment blocks nicotine-induced DA release in the nucleus accumbens [52]. However, nicotine self-administration would be rather dependent upon CB1r activation located within the VTA [56]. Conditional CB1r function on cholinergic neurons is unlikely directly modulating the reward circuit in this framework [57] but blocking CB1r alone supported its role in the hedonic aspect, sensitivity, and pursuit of reward [47, 58, 59, 60]. Blockade of CB1r also increases DA levels [61], and stimulation of DA receptors affects ACh release in the neocortex and HPC but not the striatum (the ventral part designated as the nucleus accumbens and dorsal part) [30, 62]. However, direct BF efferences that display co-localized CB1r at intra-amygdala terminal sites could regulate additional mental functions and emotional-related behaviors. The amygdala has been involved in regulating reward and motivated-related behaviors [63, 64, 65] and exhibits massive interconnections and interrelated functional relationships with the nucleus accumbens, as well as the prefrontal cortex and the ventral tegmental area, several crucial brain regions involved in motivational processing [43, 66] as well as the dorsal striatum: mounting evidence outlined the role of striatal CIN in motivational functions [67].

We also did not report significant differences in the aforementioned deletion in an effort-based choice schedule when asked decisions were based only upon pellet ratio (1 versus 3) and associated 10 lever presses for the highest reward outcome, or 1 lever press for 1 pellet delivery. All animals were able to discriminate this high reward-effort ratio that strengthened sustained motivation (as compared to control) observed in PR.

### Implications for neuropsychiatric disorders

Psychiatric disorders including schizophrenia could be manifested by short-term memory impairments [68], but common antipsychotic (APD) medication efficiency is minimal across this cognitive domain [69]. Pharmacological trials focused on studies on other drugs beyond APD [70], including cannabinoid treatments from which mechanisms of action have only begun to emerge. Our results outline promising challenging therapeutical targets regarding cannabinoid compounds activating CB1r and subsecond modulation of cholinergic release in the HPC to the extended limbic system and the PFC in particular. Further circuit manipulations using *Cre* expression on cholinergic neuronal populations exhibiting presynaptic CB1r will delineate restricted pathways and highlight local neuronal networks for cognitive processing efficiency.

## Conclusion

In conclusion, CB1r deletion on cholinergic neurons induces a predominant discrete improvement of cognitive performances represented by short-term memory abilities i.e., working memory, also uncoupled from motor action *per se* and physiological parameters such as pain sensitivity and temperature regulation. A sustained motivation was evidenced, and motor bias and temporary delay perception would be poorly involved in such reinforcing behaviors. However, CB1r deleted animals provided higher motor responses in temporally-control behavior as compared to the control mice. This array of improvement and selective processes that subserve cognition emphasizes relevant pharmacological targets and provides novel insights into understanding eCB function in both normal and pathological states.

## Acknowledgments

I would like to thank Dr. Joseph F. Cheer for his help with this work. This work was supported by the NIH.

